# Circulating senescent myeloid cells drive blood brain barrier breakdown and neurodegeneration

**DOI:** 10.1101/2023.10.10.561744

**Authors:** C. Matthias Wilk, Flurin Cathomas, Orsolya Török, Jessica Le Berichel, Matthew D. Park, George R. Heaton, Pauline Hamon, Leanna Troncoso, Brooks P. Scull, Diana Dangoor, Aymeric Silvin, Ryan Fleischmann, Meriem Belabed, Howard Lin, Elias Merad Taouli, Steffen Boettcher, Markus G. Manz, Julia K. Kofler, Zhenyu Yue, Sergio A. Lira, Florent Ginhoux, John F. Crary, Kenneth L. McClain, Jennifer L. Picarsic, Scott J. Russo, Carl E. Allen, Miriam Merad

## Abstract

Neurodegenerative diseases (ND) are characterized by progressive loss of neuronal function. Mechanisms of ND pathogenesis are incompletely understood, hampering the development of effective therapies. Langerhans cell histiocytosis (LCH) is an inflammatory neoplastic disorder caused by hematopoietic progenitors expressing MAPK activating mutations that differentiate into senescent myeloid cells that drive lesion formation. Some patients with LCH subsequently develop progressive and incurable neurodegeneration (LCH-ND). Here, we show that LCH-ND is caused by myeloid cells that are clonal with peripheral LCH cells. We discovered that circulating *BRAF*V600E^+^ myeloid cells cause the breakdown of the blood-brain barrier (BBB), enhancing migration into the brain parenchyma where they differentiate into senescent, inflammatory CD11a^+^ macrophages that accumulate in the brainstem and cerebellum. Blocking MAPK activity and senescence programs reduced parenchymal infiltration, neuroinflammation, neuronal damage and improved neurological outcome in preclinical LCH-ND. MAPK activation and senescence programs in circulating myeloid cells represent novel and targetable mechanisms of ND.

## Introduction

Neurodegenerative diseases (ND) are characterized by progressive loss of neuronal function. Mechanisms of ND pathogenesis are incompletely understood, hampering the development of effective therapies. A progressive, incurable neuroinflammatory condition (LCH-ND) arises in approximately 10% of patients with Langerhans cell histiocytosis (LCH)^1,2^. LCH is an inflammatory neoplastic disorder caused by hematopoietic progenitors expressing MAPK activating mutations that differentiate into senescent mononuclear phagocytes (MNP) that drive lesion formation^3,4^. LCH therefore represents an informative disease model to investigate mechanisms by which hematopoietic myeloid cells can induce neuroinflammation and neurodegeneration.

LCH-ND is a devastating complication of LCH patients that can arise years after the initial systemic disease is cured. Clinically, LCH-ND is characterized by progressive inflammatory lesions in the cerebellum, basal ganglia and brainstem with associated progressive neurologic clinical findings including ataxia, dysarthria and dysmetria^5^. LCH-ND was initially considered a para-neoplastic or auto-immune condition, due to the lack of CD207^+^ cells typically observed in systemic LCH lesions and the presence of T cell infiltrates in the rare cases of reported biopsies^6,7^. Optimal strategies for surveillance and treatment of LCH are poor, and the etiology of LCH-ND remains uncertain. Historically, therapeutic approaches for LCH-ND included observation, immune suppression and chemotherapy^1,8,9^.

We have previously demonstrated that multisystem LCH is caused by MAPK activating mutations, (most commonly *BRAF*V600E) that occur in early hematopoietic progenitors^10,11^. We validated *BRAF*V600E as a driver mutation that recapitulated the LCH phenotype, when expression was enforced in myeloid progenitors^10^. We further demonstrated that the *BRAF*V600E mutation induces an oncogene-driven senescence program in early hematopoietic precursors that reduces their proliferation potential. In these cells, survival is supported by expression of anti-apoptotic molecules such as Bcl-xL and senescence-associated secretory proteins (SASP) that skew the differentiation of *BRAF*V600E^+^ hematopoietic progenitors into senescent MNP that seed peripheral tissues^3^. Senescent MNP that accumulate in tissues drive the formation of inflammatory and granulomatous LCH lesions, leading to tissue damage. Notably, therapies that block SASP and dis-inhibit resistance to cell death decrease disease burden and prolong survival in mouse models^3^.

The relationship between systemic LCH and LCH-ND is not well defined. There are associations between the risk of developing LCH-ND and disseminated LCH, progressive or relapsed systemic disease, *BRAF*V600E mutation, and lesions in the skull base, the pituitary and brain parenchyma^5^. Importantly, we recently identified circulating *BRAF*V600E^+^ peripheral blood mononuclear cells (PBMC) in the blood of patients with LCH-ND, even in the absence of active systemic lesions. We also detected *BRAF*V600E^+^ myeloid cells in brain biopsies of patients with LCH-ND^3,12^ and demonstrated that, similar to peripheral LCH cells, pathogenic MNP that accumulate in the brain of human LCH-ND exhibit features of senescence^3^. Based on these observations and the clinical patterns of LCH-ND, we hypothesized that LCH-ND arises from hematopoietic precursors that are clonal with peripheral LCH lesions and that senescence programs in infiltrating myeloid cells may also contribute to brain injury in patients with LCH-ND.

Taking advantage of the LCH model of persistent MAPK activation in hematopoietic cells, we sought to define mechanisms by which MAPK activation in MNP could drive neuroinflammation and neurodegeneration. Resident and recruited MNP populations in the central nervous system (CNS) are increasingly recognized to play central roles in a wide range of neurodegenerative conditions^13^ by orchestrating immune cell activation. This study highlights a novel mechanism for monocyte-derived CD11a^+^ macrophages to induce CNS neurodegenerative diseases that can be reversed with MAPK inhibition and senolytic therapy.

## Results

### Genetic fate-mapping reveals that circulating *BRAF*V600E^+^ myeloid cells accumulate in the brains of mice with LCH-like disease

To study the potential cellular determinants of LCH-ND, we designed transgenic mice that express *BRAF*V600E, the most common somatic mutation in LCH, at different stages of hematopoietic differentiation, recapitulating key features of LCH^3,10^. Specifically, we found that multi-system LCH developed in mice, in which we enforced the *BRAF*V600E mutation under the promoter of the *Scl* gene that is expressed in long-term and short-term hematopoietic stem cells (HSC) in the bone marrow and showed that *BRAF*V600E^+^ MNP were the main drivers of tissue injury in the periphery^3^.

To explore whether LCH-ND was also driven by circulating *BRAF*V600E^+^ cells, we used the above model of LCH models in which a conditional *BRAF*V600E mutation was enforced under the promoter of the *Scl* gene (**Fig. 1a**). As previously described, we genetically engineered somatic mosaicism for a *BRAF*V600E allele linked to a yellow fluorescent protein (YFP) in HSC using tamoxifen (Tam)-inducible targeting in *Scl*^creERT^ mice, which we named *BRAF*V600E*^Scl^*and used *BRAF* wild-type (*BRAF*wt*^Scl^*) as control littermates. Thus, YFP-expressing cells carried the *BRAF*V600E mutation in *BRAF*V600E*^Scl^* mice, whereas YFP-expressing cells in *BRAF*wt*^Scl^* animals did not carry the mutation but underwent Rosa26-locus driven Cre recombination. Strikingly, in addition to systemic LCH lesions, we also detected YFP-reporter-tagged, *BRAF*V600E-mutated cells in the brains of *BRAF*V600E*^Scl^* animals between week 8 and 12 after tamoxifen-induced cre-recombination (**Fig. 1b**). We further characterized these cells as CD11a^+^ macrophages, a monocyte-derived population recently described in the CNS vasculature as having the potential to infiltrate brain parenchyma^14^. The population of CD11a^+^ macrophages was the only myeloid cell population expressing the *BRAF*V600E mutation in the brains of *BRAF*V600E*^Scl^*mice. This CD11a^+^ macrophage population was also enriched up to 50-fold compared to control animals. *BRAF*V600E^+^ cells were notably not detected in either microglia or perivascular macrophages (PVM) (**Fig. 1b**). The percentage of YFP-tagged cells in the brains of mice correlated with the percentage of reporter-tagged cells in the peripheral blood and was significantly higher in *BRAF*V600E*^Scl^* animals compared to *BRAF*wt*^Scl^* animals (**Fig. 1c**).

**Figure 1:**
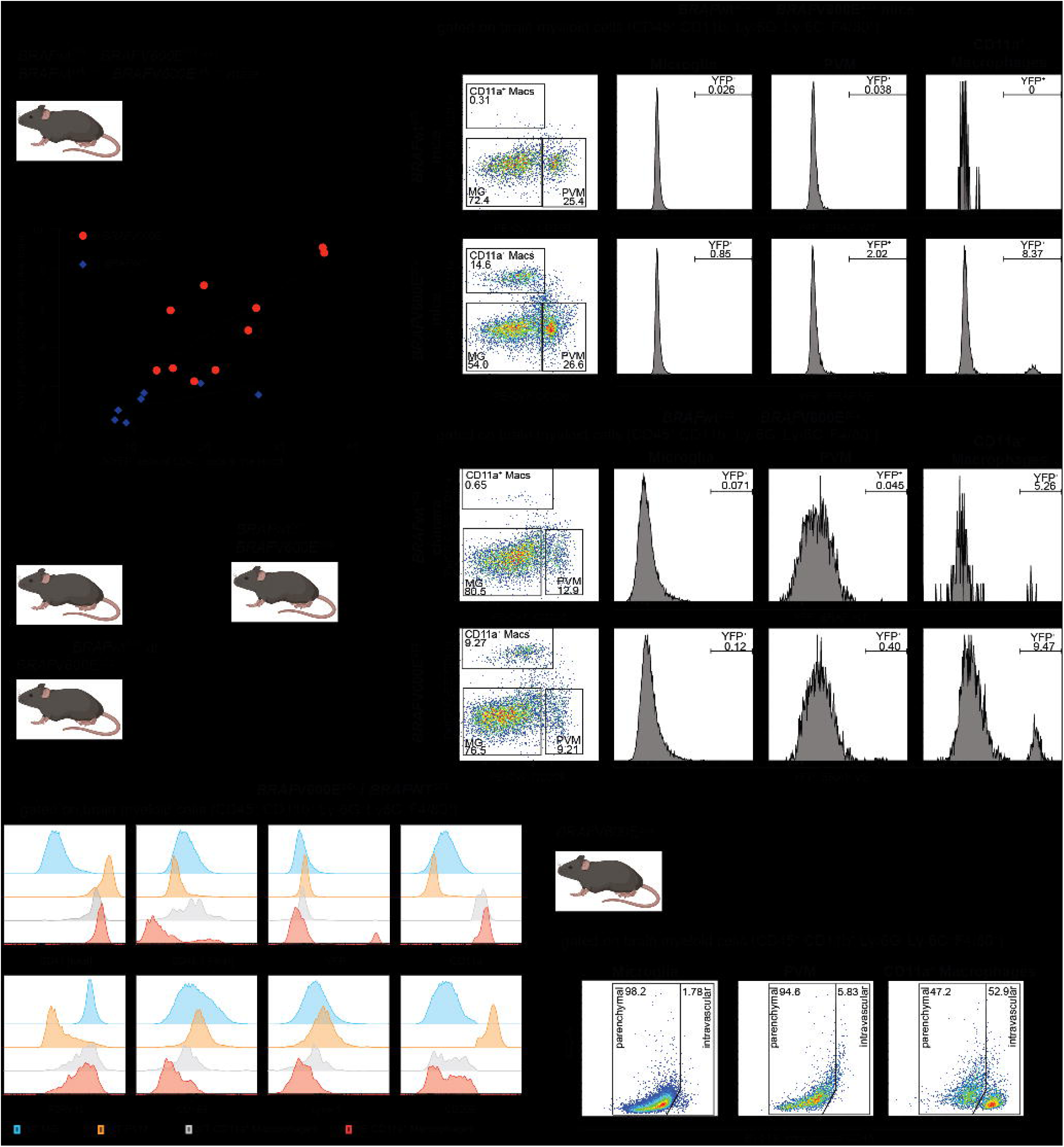
Genetic fate-mapping reveals that circulating *BRAF*V600E^+^ myeloid cells accumulate in the brains of mice with LCH-like disease. **a**, experimental setup of *Scl* and *Map17*-based LCH mouse models: Tamoxifen was applied for Cre recombination and mice were then terminally analyzed after 8-16 weeks as indicated. **b,** representative spectral cytometry pseudocolor and histogram plots of microglia (MG), perivascular macrophages (PVM) and CD11a^+^ macrophages from *BRAF*V600E*^Scl^*mice and *BRAF*wt*^Scl^* control mice. The abundance of reporter-tagged cells in these populations is displayed in the respective histogram plots. **c,** correlation between percentage of YFP-tagged cells in the peripheral blood and percentage of YFP-tagged cells in the brains of *BRAF*V600E*^Scl^*mice and *BRAF*wt*^Scl^* control mice (n=7-10). **d,** experimental setup to generate *BRAF*V600E*^Scl^* chimera and *BRAF*wt*^Scl^*control chimera. **e,** representative spectral cytometry pseudocolor and histogram plots of the brain myeloid cell populations of *BRAF*V600E*^Scl^*chimera and *BRAF*wt*^Scl^* control chimera. **f,** expression of key lineage markers in the different brain myeloid cell populations of *BRAF*V600E*^Scl^*chimera and *BRAF*wt*^Scl^* control chimera. **g,** experimental setup of intravascular cell staining to determine vascular vs parenchymal localization of cells: *BRAF*V600E*^Scl^* mice were injected with a BV510-conjugated anti-CD45 antibody and were shortly after terminally analyzed (n=4 mice). **h,** Representative spectral cytometry pseudocolor plots of intravenous CD45 staining for each myeloid cell population.

To confirm whether *BRAF*V600E^+^ macrophages that accumulated in the brain of *BRAF*V600E*^Scl^* animals arose from circulating precursors and not locally, we developed another model of MAPK activating mutations expressed exclusively in bone marrow HSC by enforcing *BRAF*V600E mutation under the *Map17* gene promoter known to be expressed in long-term bone marrow HSC, which we refer to as *BRAF*V600E*^Map17^* mice (**Fig. 1A**)^15^. Similar to the *BRAF*V600E*^Scl^* model, *BRAF*V600E*^Map17^*mice developed systemic LCH lesions (**Fig. S1a-b**), again emphasizing that multisystem LCH is a myeloid inflammatory neoplasia driven by MAPK activating mutation expressed in early HSC^4^. Importantly and similar to *BRAF*V600E*^Scl^* mice, we detected a large population of *BRAF*V600E^+^ CD11a^+^ macrophages in the brain parenchyma of *BRAF*V600E*^Map17^* mice between week 12 and 16 after tamoxifen-induced Cre recombination (**Fig. S1c-d**), suggesting that circulating *BRAF*V600E^+^ myeloid cells indeed give rise to *BRAF*V600E^+^ CD11a^+^ macrophages.

To further ensure that *BRAF*V600E^+^ CD11a^+^ macrophages which accumulated in the brain of these mice derived from circulating mutated cells and did not arise locally, we generated bone marrow chimeric animals (CD45.2 *BRAF*V600E*^Scl^*:CD45.1 wild-type) animals in which bone marrow cells isolated from *BRAF*V600E*^Scl^*or lineage tracing control mice (*BRAF*wt*^Scl^*) were injected into CD45.1^+^ congenic host mice (**Fig. 1d**). In order to prevent damage to the blood brain barrier (BBB) induced by pre-transplant conditioning that could artifactually promote translocation of circulating cells into the brain parenchyma, host mice were irradiated with a lead head-shield, as previously described^16^. We found that similar to transgenic mice, chimeric mice also developed systemic LCH lesions (**Fig. S1e-f**) as well as brain lesions with accumulation of *BRAF*V600E CD11a^+^ macrophages (**Fig. 1e, Fig. S1e-f**). Of note, chimeric mice transplanted with BM from *BRAF*wt*^Scl^*and *BRAF*V600E*^Scl^* donor mice did not differ in terms of donor chimerism (**Fig. S1g**). CD11a^+^ macrophages originated from the donor bone marrow (CD45.2^+^) and expressed high levels of the cell type-defining integrin alpha L (CD11a) but lacked the expression of markers of PVM (e.g., CD169, LYVE-1, CD206) and showed modest induction of tissue resident microglia markers (e.g., P2RY12) (**Fig. 1f**), confirming that these cells do not present cues of local resident origin.

We next sought to determine whether CD11a^+^ macrophages remained restricted to the vasculature or had the potential to invade the brain parenchyma. We intravenously injected *BRAF*V600E*^Scl^* mice with an anti-CD45 antibody conjugated to a fluorescent dye to tag intravascular cells prior to brain isolation and analysis (**Fig. 1g**). Nearly 50% of brain associated CD11a^+^ macrophages were unstained, suggesting that a substantial fraction of the CD11a^+^ macrophages while derived from the blood circulation, had extravasated into the brain parenchyma of *BRAF*V600E*^Scl^*mice (**Fig. 1h**).

### LCH mice with brain-infiltrating *BRAF*V600E-mutated cells share phenotypic and transcriptional characteristics with human LCH-ND

In humans, imaging studies and clinical features typically localize LCH lesions and CNS injury to the brainstem (pons, medulla oblongata) and cerebellum as well as to areas with limited BBB (pituitary, vermis, choroid plexus)^5^. We recently reported enrichment of *BRAF*V600E^+^ cells in brainstem and cerebellum with minimal infiltration in the temporal and frontal lobes of a whole brain autopsy from a young man who died from progressive LCH-ND^17^ (**Fig. 2a-b**). Remarkably, the pattern of neurodegeneration in human LCH-ND matched the main sites of parenchymal infiltration in the *BRAF*V600E*^Scl^* chimera (**Fig. 2c-d**). Of note, the infiltration pattern in our *BRAF*V600E*^Scl^* chimera was diffuse with a lack of granulomatous lesions (**Fig. 2e**, right). By comparison, reporter-tagged cells were virtually absent in brain tissue from the control chimera (**Fig. 2e**, left).

**Figure 2:**
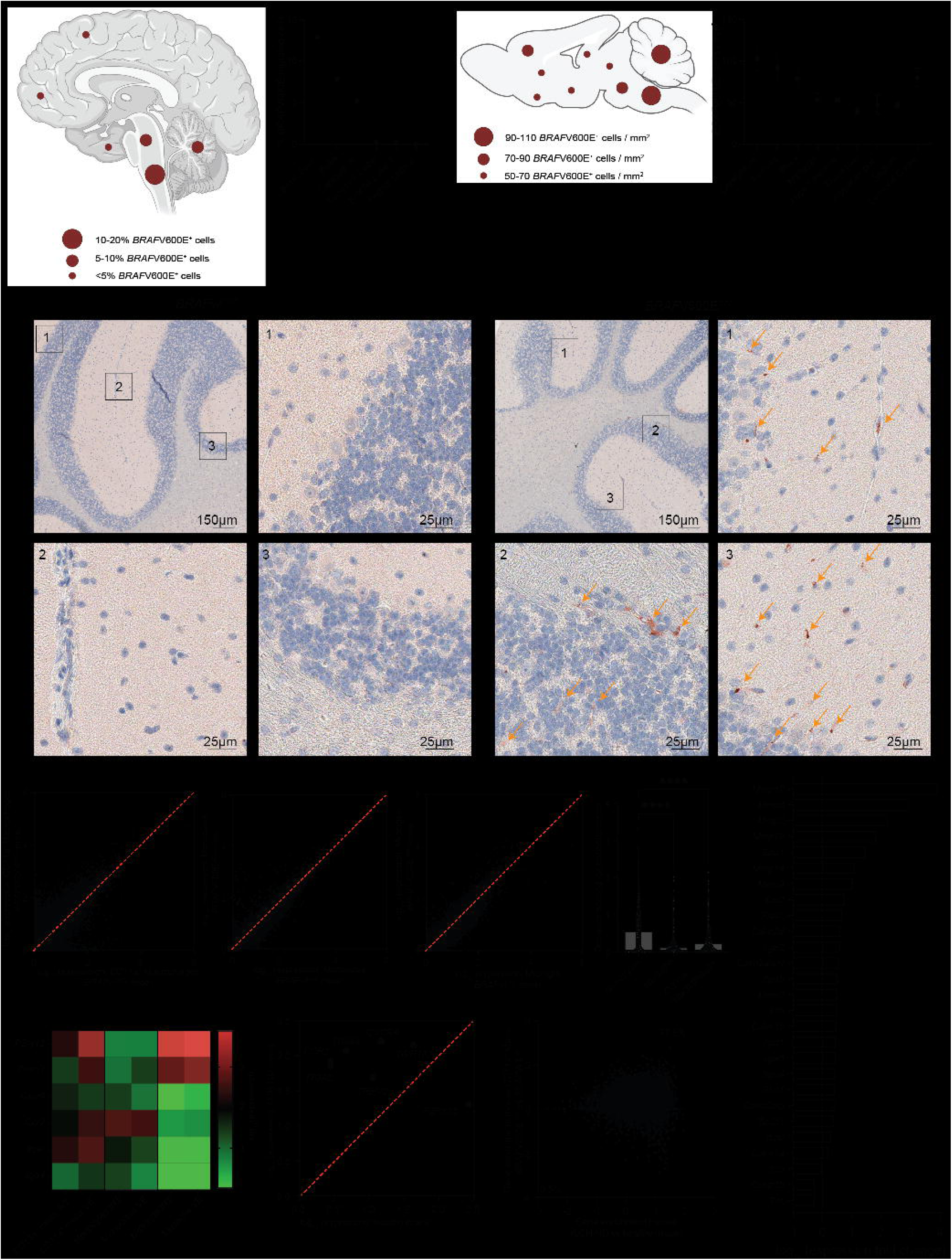
LCH mice with brain-infiltrating *BRAF*V600E-mutated cells share phenotypic and transcriptional characteristics with human LCH-ND. **a,** manifestation of LCH-ND in a schematic human brain and **b,** quantification of *BRAF*V600E transcripts from this specimen. **c,** visualization of the extent to which mouse brains are affected by infiltration with *BRAF*V600E-mutated cells based on the quantification from different brain regions depicted in **d** (n=4 mice). **e,** Representative IHC images from the brain of a *BRAF*wt*^Scl^*chimera on the left and *BRAF*V600E*^Scl^*chimera on the right stained for YFP-tagged, BM-derived cells marked with yellow arrows. In *BRAF*wt*^Scl^* chimera, these cells are lineage-traced unmutated cells (left) and in *BRAF*V600E*^Scl^* chimera, these stained cells are *BRAF*V600E^+^ cells (right). **f-k**, CD11a^+^ macrophages (**f**), peripheral blood monocytes (**g**) as well as microglia (**h**) from *BRAF*wt*^Scl^* and *BRAF*V600E*^Scl^* mice were subjected to bulk RNA sequencing from each 10 mice per group (pooled). Expression profiles of CD11a^+^ macrophages (**f**), monocytes (**g**) and microglia (**h**) from *BRAF*wt*^Scl^* mice were compared with those from *BRAF*V600E*^Scl^* mice in **i** by analysis of the deviance of transcriptomes showing that the transcriptomic remodeling was strongest in CD11a^+^ macrophages. While monocytes and CD11a^+^ macrophages in *BRAF*V600E*^Scl^* mice are *BRAF*V600E mutated, microglia is unmutated in *BRAF*V600E*^Scl^* mice. **j**, the transcriptional remodeling in CD11a^+^ macrophages was driven by senescence-associated genes as well as Matrix Metalloproteinases and integrins. **k**, comparative analysis of gene expression of key lineage markers in mouse CD11a^+^ macrophages, peripheral blood monocytes and microglia. **l,** comparison of these lineage markers between a human bulk RNA sequencing data set from an LCH-ND case and human myeloid cells from a healthy brain demonstrating an enrichment for CD11a^+^ macrophage-defining *Itgal* (encoding integrin alpha L chain, CD11a) as well as *Itga4* (encoding integrin alpha 4 chain, CD49d). **m,** cross-species comparison of mouse *BRAF*V600E^+^ CD11a^+^ macrophages and human LCH-ND material showing that 57.5% of the genes were enriched in mouse and human LCH material. Data in **d** are shown as means ± s.e.m, ****p<0.0001. Abbreviations: LCH-ND: neurodegenerative LCH, wt: wild-type, VE: *BRAF*V600E

To further establish the pathogenic role of blood-derived *BRAF*V600E^+^ CD11a^+^ macrophages, we profiled purified CD11a^+^ macrophages, peripheral blood monocytes and microglia from *BRAF*V600E*^Scl^*and *BRAF*wt*^Scl^* mice using bulk RNA sequencing (bulk RNAseq). In comparing the mRNA transcriptome profiles of subsets of each myeloid cell subset from *BRAF*wt*^Scl^* and *BRAF*V600E*^Scl^*mice (**Fig. 2f-h**), we generated a deviance score to represent the divergence of gene expression driven by the *BRAF*V600E mutation, wherein a score of 0 would indicate complete concordance between genotypes (**Fig. 2i**). We found that the transcriptomic differences observed in mutant versus wild-type CD11a^+^ macrophages were the most dramatic, compared to that computed for *BRAF*V600E^+^ and wild-type monocytes and microglia (**Fig. 2i**), suggesting that the induction of the MAPK pathway activation by *BRAF*V600E significantly impacts the molecular phenotype of the CD11a^+^ macrophages, in line with the anticipated effect of Cre recombination in *BRAF*V600E*^Scl^* mice. Of note, while monocytes from *BRAF*V600E*^Scl^*mice harbor the *BRAF*V600E mutation, microglia from *BRAF*V600E*^Scl^* mice does not.

We had previously shown that an oncogene-induced senescence program is the driver of inflammation in CD207^+^ cells that infiltrate systemic LCH lesions^3^. Thus, we measured the expression of canonical senescence genes in wild-type and *BRAF*V600E CD11a^+^ macrophages and confirmed upregulation of senescence-related genes in *BRAF*V600E*^Scl^* CD11a^+^ macrophages, compared to wild-type cells. Among these upregulated genes, we identified cell cycle control genes encoding the anti-apoptotic Bcl-2 family member Bcl-xL (*Bcl2l1*), uPAR (*Plaur*), and matrix metalloproteinases, which are all classically upregulated during oncogene-induced senescence (**Fig. 2j**). In addition, *BRAF*V600E^+^ CD11a^+^ macrophages overexpressed several integrin sub-units in addition to αL (CD11a, *Itgal*) including subunits α4 (*Itga4*), α5 (*Itga5*), α6 (*Itga6*) and β1 (*Itgb1*), suggesting a potential role for this integrin program in mediating infiltration of brain tissue by *BRAF*V600E^+^ CD11a^+^ macrophages (**Fig. 2k**). CXCR4 was expressed on CD11a^+^ macrophages, in line with our observation that these are bone marrow-derived cells (**Fig. 2k**). We also compared this transcriptional dataset to a case of LCH-ND, for which an RNAseq data set is publicly available^18^. Consistent with our observations in mice, we found that mRNA expression of the lineage genes depicted in **Fig 2k** was more enriched in human LCH-ND, than in bulk profiling of healthy human brain myeloid cells (**Fig. 2l**). In a more comprehensive analysis, we generated a shared mRNA library between our murine bulk RNAseq dataset from CD11a^+^ macrophages from *BRAF*V600E*^Scl^*mice with the published dataset of healthy and LCH-afflicted brain tissue – akin to published strategies applied for single-cell transcriptomic datasets^19,20^. We compared the expression patterns of genes that are enriched in murine *BRAF*V600E^+^ CD11a^+^ macrophages, compared to their wild-type counterparts, with those differentially expressed between human LCH-ND and healthy human brain tissue. From this homology analysis, we found that nearly 60% of genes enriched in *BRAF*V600E^+^ CD11a^+^ macrophages are also more highly expressed in human LCH-ND, indicating significant concordance between the transcriptional activity between human LCH-ND and brain infiltration by CD11a^+^ macrophages in our mouse model (**Fig. 2m**), supporting the likely contribution of the gene program of CD11a^+^ macrophages in human LCH-ND, analogous to its role in our murine model of LCH-ND.

### *BRAF*V600E*^Scl^* chimera have a compromised BBB and exhibit behavioral and neurologic abnormalities

Systemic inflammation has been shown to alter the BBB and promote the extravasation of circulating myeloid cells into the brain^13^. Because senescence and systemic inflammation are hallmark features of LCH, we sought to examine whether BBB is altered in mice with multisystem LCH. Consistent with patterns of systemic inflammation in LCH, we found elevated pro-inflammatory cytokines in the brains of *BRAF*V600E*^Scl^* chimera, as compared to their wild-type counterparts (**Fig. 3a**, **b**). To measure if systemic inflammation was associated with increased BBB leakage in our model, we intravenously injected LCH animals with the albumin-bound Evans blue dye and measured its extravasation into the brain parenchyma. We found significantly increased dissemination of Evans blue dye throughout the brains of LCH mice, compared to that of control animals, indicative of an altered BBB that enabled the translocation of small molecules (**Fig.3** **c, d**). To directly measure an active translocation of blood-derived cytokines, we intravenously injected *BRAF*V600E*^Scl^* and *BRAF*wt*^Scl^*control animals with a biotinylated version of the inflammatory cytokine interleukin 1 beta (IL-1β) in *BRAF*V600E*^Scl^* mice and *BRAF*wt*^Scl^*control animals. We used an avidin-bound fluorescent dye to measure the accumulation of translocated IL-1β in the brain parenchyma of injected mice. Importantly, we found a significant increase of extravasated IL-1β in the brain of LCH mice compared to control mice establishing the increased permeability of the BBB in LCH mice (Fig. 3e-g).

**Figure 3:**
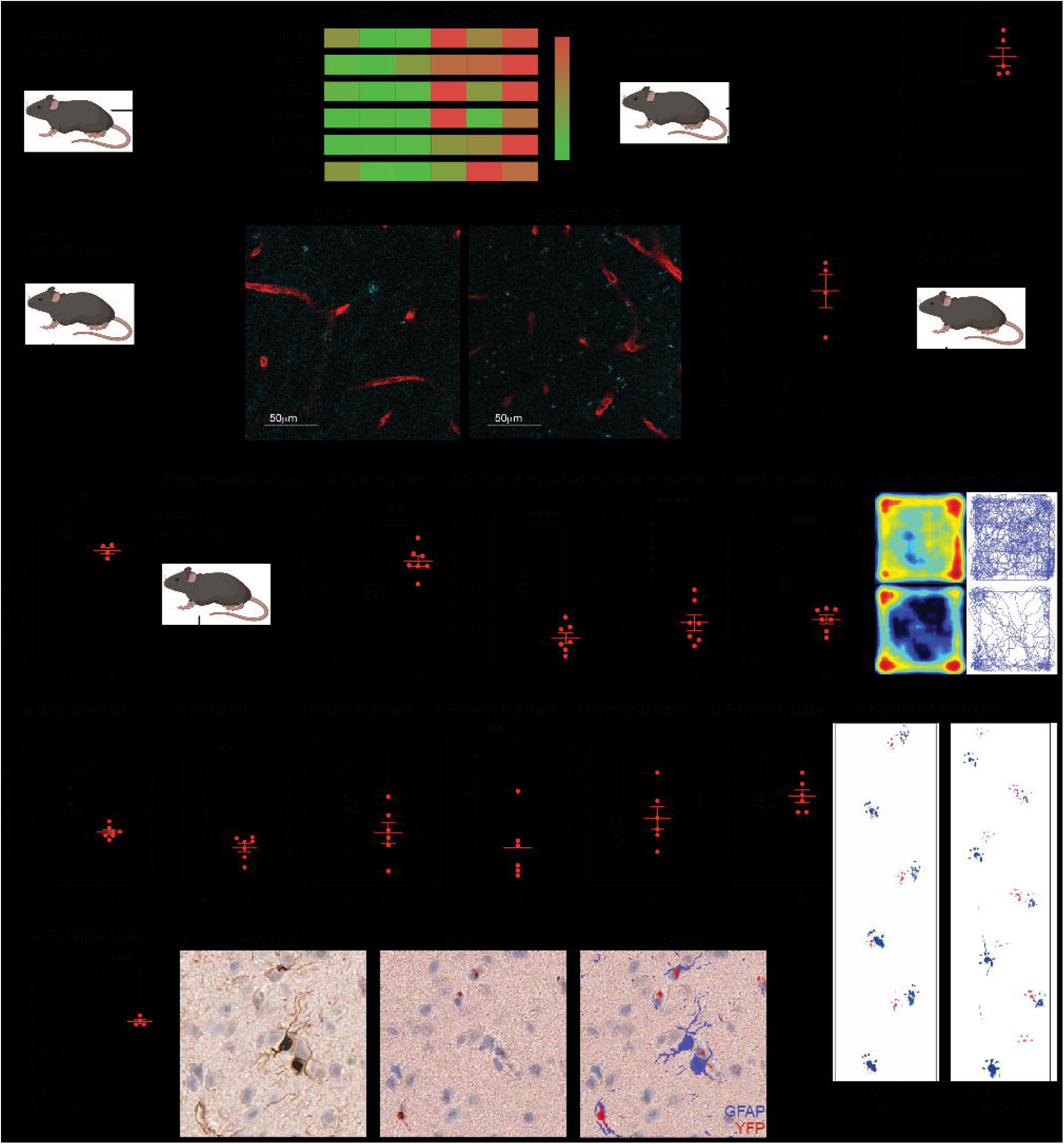
*BRAF*V600E*^Scl^*chimera have a compromised BBB and exhibit behavioral and neurologic abnormalities. **a,** experimental setup to quantify inflammatory cytokines from brain lysates from *BRAF*V600E*^Scl^* and *BRAF*wt*^Scl^*mice. **b**, multiplexed cytokine detection in the brains of *BRAF*V600E*^Scl^*and *BRAF*wt*^Scl^* control mice (n=3 mice per group). **c**, experimental setup of an Evans Blue-based assay to quantify the permeability of the BBB for small molecules showing increased permeability in *BRAF*V600E*^Scl^*mice (**d**, n=4-5 mice per group). **e**, experimental setup to assess and quantify the transition of biotinylated IL-1β into the brain of *BRAF*V600E*^Scl^* mice and *BRAF*wt*^Scl^* control mice. **f**, representative images of IL-1β (cyan) and Collagen IV (red) showing a scattered perivascular signal derived from biotinylated IL-1β that can be detected in *BRAF*V600E*^Scl^*mice but not *BRAF*wt*^Scl^* control mice (n=3-4 mice per group). **g**, quantification of IL-1β-derived fluorescent signal shows a significant increase in extravasation of biotinylated IL-1β into the brain parenchyma in *BRAF*V600E*^Scl^* mice compared to *BRAF*wt*^Scl^*mice. **h**, experimental setup to assess the pericyte coverage of brain vessels from *BRAF*V600E*^Scl^* and *BRAF*wt*^Scl^*chimera. **i**, *BRAF*V600E*^Scl^* chimera have a decreased blood vessel coverage with pericytes compared to *BRAF*wt*^Scl^*chimera (n=3-4 mice per group). **j**, experimental setup for behavioral assays and histological studies for **k-x**. **k-o**, performance of *BRAF*V600E*^Scl^*chimera (VE) and *BRAF*wt*^Scl^* chimera (WT) in an open field setup showing a deteriorated performance with reduced resting time (**k**), distance traveled (**l**), time in center (**m**) and median velocity (**n**) summarized in the activity plots, **o** (n=7 mice per group). Grip strength (**p**) and latency to fall (**q**) quantification of *BRAF*V600E*^Scl^* and *BRAF*wt*^Scl^*chimera in a Rotarod assay (n=7 mice per group). **r-v**, footprint analysis of Hindlimb and Forelimb Stride **(r, s)** as well as Hindlimb and Forelimb base as signs of a motor deficit in *BRAF*V600E*^Scl^* chimera (**t, u**, n=6-10 mice per group). **v,** representative case of unilateral paralysis in a *BRAF*V600E*^Scl^* chimeric mouse compared to a *BRAF*wt*^Scl^*mouse; hind paws painted with blue and front paws painted with red ink. **w,** quantification of Purkinje cells in *BRAF*V600E*^Scl^* chimera and *BRAF*wt*^Scl^*chimera (n=4 mice per group). **x**, representative images of multiplex immunohistochemistry of the brains of *BRAF*V600E*^Scl^* chimera staining for activated astrocytes (anti-GFAP, left), *BRAF*V600E-mutated cells (anti-YFP, middle) and showing co-localization (overlay, right). *p<0.05, **p<0.01, ***p<0.001, ****p<0.0001, n.s. = not significant. Data are shown as means ± s.e.m. Abbreviations: r.o.: retro orbital, GFAP: Glial Fibrillary Acidic Protein.

To further establish the breakdown of BBB in LCH mice, we measured the pericyte coverage of blood vessels in the brains of *BRAF*V600E*^Scl^* and *BRAF*wt*^Scl^*chimera as a surrogate of BBB integrity (**Fig. 3h**). Accordingly, we documented significantly reduced pericyte coverage of the BBB of *BRAF*V600E*^Scl^*chimera relative to *BRAF*wt*^Scl^* chimera, indicating a breakdown of the BBB that likely facilitated the extravasation of inflammatory cytokines such as IL-1β and the migration of bone marrow-derived *BRAF*V600E mutated cells into the brain parenchyma (**Fig. 3i**).

To assess the pathogenic consequence of the BBB breakdown and increased extravasation of inflammatory cytokines and myeloid cells in the brain parenchyma of LCH mice, we measured the behavioral and neurological phenotypes of *BRAF*V600E*^Scl^* chimera using a range of behavioral tests (**Fig. 3j**). We evaluated murine behavior in an Open Field Assay and found that *BRAF*V600E*^Scl^* chimera performed significantly worse compared to control chimera in most relevant parameters such as increase in resting time (**Fig. 3k**), decreased total distance traveled (**Fig. 3l**) and reduced time in center and a lower median velocity (**Fig. 3m-o**). Using a rotarod assay, we found that *BRAF*V600E*^Scl^*chimera have reduced grip strength and a shorter latency to fall compared to WT mice (**Fig. 3p, q**), suggesting profound motor deficit in LCH mice. Because CD11a^+^ macrophages mostly accumulated in the cerebellum, we further focused on motor function assays. Using a footprint assay, we observed a significantly shortened hindlimb and forelimb stride length (**Fig. 3r, s**) and a trend towards a broader hind- and forelimb base (**Fig. 3t, u**). Sporadic cases of unilateral paralysis had also been observed (**Fig. 3v**). Using multiplex immunohistochemistry staining, we also measured Purkinje damage and found a significant loss of Purkinje cells and astrocyte activation in close proximity to *BRAF*V600E-mutated cells in the brain of *BRAF*V600E*^Scl^*chimera in line with the behavioral and neurological phenotypes observed in these mice (**Fig. 3w, x**).

### Accumulation of circulating, *BRAF*V600E^+^ cells is a driver of LCH-like disease and combined senolytic / MAPK inhibitor therapy alleviates the disease burden

Bcl-xL is highly expressed in systemic LCH lesions, and we have previously shown that treatment of *BRAF*V600E*^Scl^* mice with the senolytic Bcl-xL inhibitor can deplete LCH cells^3,11^. We therefore tested the ability of MAPK pathway inhibition (trametinib [T], MEK inhibitor) and senolytic therapy (navitoclax [N], Bcl-xL inhibitor) to reduce the accumulation of mutant CD11a^+^ macrophages in the brain and the associated neuroinflammation (**Fig. 4a, d**). MAPK pathway inhibition monotherapy has become a mainstay in relapsed and refractory LCH, but fails to eradicate the LCH clone in mice and humans; it is also less effective in LCH-ND, compared to systemic LCH^21,22^. In this study, combining MEK inhibition (T) and Bcl-xL inhibition (N) led to a substantial reduction of LCH cells in the lungs of mice (**Fig. 4b, Figure S3a**). Importantly, we also found that MEK inhibition and Bcl-xL inhibition (NT) led to a substantial reduction of CD11a^+^ macrophages in the brain parenchyma mice of transgenic *BRAF*V600E*^Scl^*mice (**Fig. 4c**). We confirmed a beneficial effect of the NT combination therapy in the *BRAF*V600E*^Scl^*chimera model with a significant reduction of pathologically increased liver and spleen weight, which become infiltrated with mutant cells in disseminated LCH (**Fig. 4d-f**). Strikingly, treatment with the NT combination also corrected the behavioral impairment observed in LCH mice with a normalized performance in most aspects of the Open Field Assay (**Fig. 4g-j** and **k,** top and middle rows) which correlated with reduced accumulation of mutant cells in the brains of these mice (**Fig. 4k**, bottom row).

**Figure 4:**
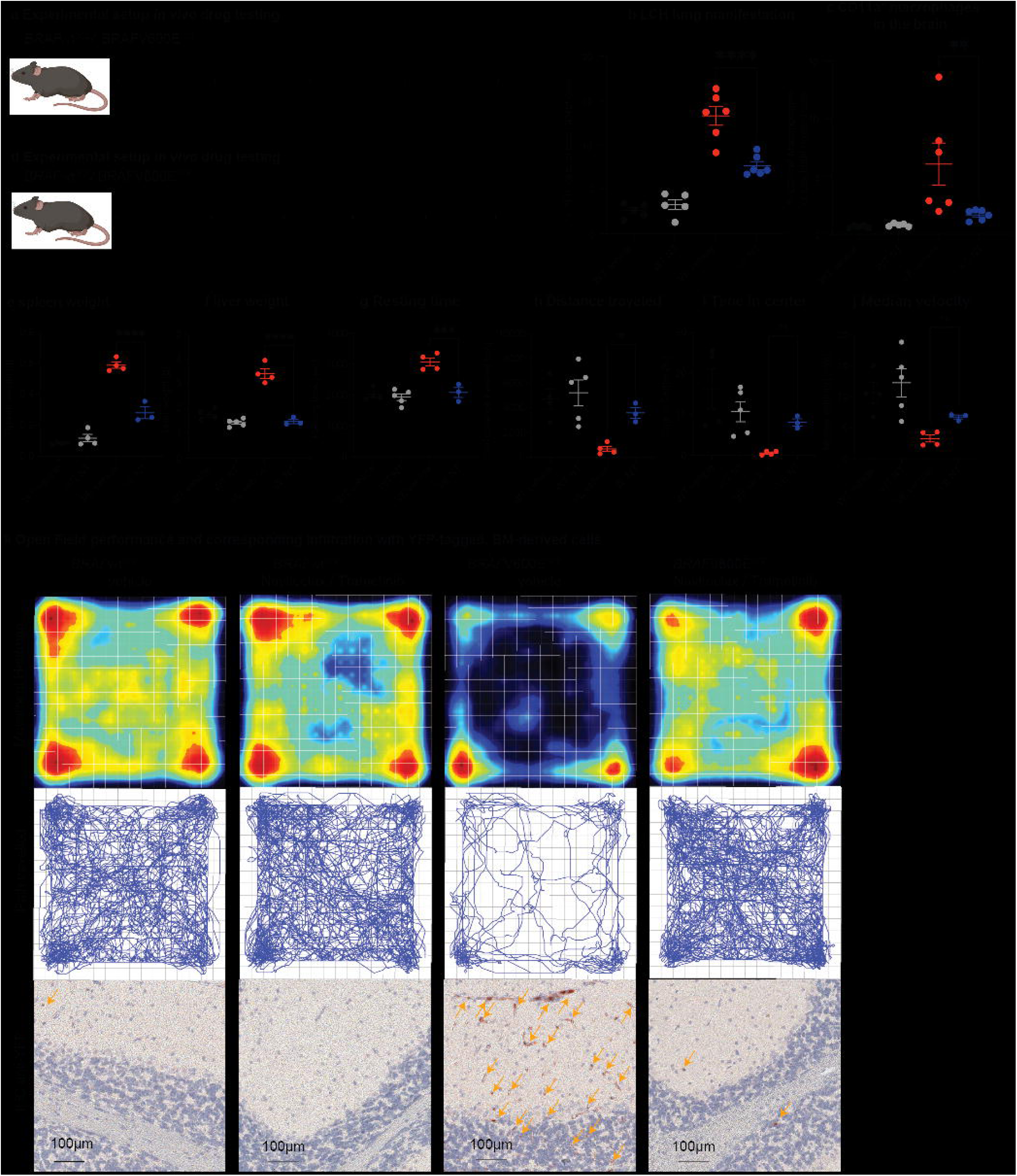
Accumulation of circulating, *BRAF*V600E^+^ cells is a driver of LCH-like disease and combined senolytic / MAPK inhibitor therapy alleviates the disease burden. **a,** experimental setup for *in vivo* drug treatment of *BRAF*wt*^Scl^*mice and *BRAF*V600E*^Scl^* mice with Navitoclax/Trametinib (NT) or vehicle control. **b,** percentage of YFP^+^ cells in the lungs of *BRAF*wt*^Scl^* (WT) and *BRAF*V600E*^Scl^*(VE) mice receiving vehicle or Navitoclax/Trametinib (NT) treatment analyzed by spectral cytometry (n= 5-6 per group). **c**, percentage of CD11a^+^ macrophages in the brains of *BRAF*wt*^Scl^* (WT) and *BRAF*V600E*^Scl^* (VE) mice receiving vehicle or NT combination treatment (n=5-6 per group). **d**, experimental setup for *in vivo* drug treatment of *BRAF*wt*^Scl^* chimera and *BRAF*V600E*^Scl^*chimera with Navitoclax/Trametinib (NT) or vehicle control. **e-f,** quantification of the weight of the spleens (**e**) and the livers (**f**) of *BRAF*wt*^Scl^*(WT) and *BRAF*V600E*^Scl^* (VE) chimera receiving vehicle or NT treatment (n=3-4 mice per group). **g-j** Open Field assessment of vehicle- or NT-treated *BRAF*wt*^Scl^* (WT) and *BRAF*V600E*^Scl^* (VE) chimera with quantification of the resting time **(g)**, distance travelled **(h)**, time in the center **(i)** and mouse median velocity. **k**, movement heatmap (top row) and path traveled (middle row) in the Open Field assessment with paired anti-YFP IHC from cerebellar regions (bottom row) of the respective mice. *p<0.05, **p<0.01, ***p<0.001, ****p<0.0001, n.s. = not significant. Data are shown as means ± s.e.m. Abbreviations: NT: Navitoclax/Trametinib, IHC: Immunohistochemistry

## Discussion

LCH-ND is a devastating condition that arises in 10% of patients following the development of systemic LCH. In the most severe cases, patients develop progressive dysarthria, dysmetria, and ataxia that causes significant morbidity and can lead to death^5^. The etiology of LCH-ND has been historically poorly defined. In the past decades, the few brain biopsies studied showed inflammatory cells, but lacked CD207^+^ cells characteristic of systemic LCH lesions and granulomatous brain lesions^6,7^. LCH-ND was therefore considered a paraneoplastic or autoimmune phenomenon and typically treated with immunomodulatory therapies (e.g., intravenous immune globulin, IVIG). Importantly, we identified *BRAF*V600E^+^ cells in the brain biopsies of LCH-ND^17^. In fact, in a brain autopsy from a young patient with fatal LCH-ND, we found that >10% of cells in brainstem and cerebellum were *BRAF*V600E^+^, and anatomic concentration of *BRAF*V600E^+^ cells mirrored distribution of disease identified by T2 hyperintensity on MRI. Further, histology of LCH-ND biopsy and autopsy specimens characterized the perivascular concentration of *BRAF*V600E^+^ cells. We also identified *BRAF*V600E^+^ cells in peripheral blood of patients with LCH-ND (without systemic lesions), but did not identify persistent *BRAF*V600E^+^ cells in peripheral blood of patients cured of systemic LCH and without LCH-ND^17^ (**Fig. 2b**). Similarly, Mass et al. reported the presence of *BRAF*V600E^+^ cells in brain biopsies from patients with LCH and related disorder Erdheim-Chester Disease (ECD)^23^. These findings strongly support a clonal relationship between LCH-ND and systemic LCH. Based on our observations of the presence of circulating *BRAF*V600E^+^ cells in patients with LCH-ND and perivascular distribution of *BRAF*V600E^+^ in LCH-ND biopsies, we hypothesized that LCH-ND is caused by circulating, bone marrow-derived myeloid cells clonal with systemic LCH lesion MNPs^17^.

To define the mechanisms of LCH-ND, we evaluated three different mouse models for LCH (*BRAF*V600E*^Scl^*, *BRAF*V600E*^Map17^*, *BRAF*V600E*^Scl^* chimera) in which *BRAF*V600E is post-natally enforced in HSCs. All of these mice developed aggressive systemic LCH-like disease with risk-organ involvement. We found that circulating *BRAF*V600E^+^ cells infiltrated the brain parenchyma in late-stage disease in all three models, similar to the paradigm in human disease. LCH-ND is typically a late complication of multi-system LCH and typically develops years to decades after the systemic disease has been treated. Recapitulating this sequential pattern and long latency in mouse models is difficult. The advantage and likewise drawback, however, of transgenic mouse models is that the disease burden reflected by the number of circulating cells is higher. In our *BRAF*V600E*^Scl^* chimera, roughly 10-20% of all peripheral blood cells expresses the *BRAF*V600E mutation (**Fig 1c**), whereas the burden of circulating cells in human disease varies but is usually in the range of low single-digit percentage, mostly <5%^17^. The higher number of circulating *BRAF*V600E^+^ cells in *BRAF*V600E*^Scl^* mice and *BRAF*V600E*^Scl^* chimera then allows a more rapid onset of CNS disease that can be studied before the animals succumb to systemic LCH.

We found that human and murine brain manifestations in LCH share similar characteristics which include 1) a similar distribution pattern of inflammatory cells in the brain parenchyma 2) the presence of circulating *BRAF*V600E^+^ myeloid cells expressing a senescence transcriptional program, 3) the presence of inflammatory mediators in the brain parenchyma 4) neurological impairment that affects the cerebellum as well as the brain stem. This leads to motor symptoms such as dysdiadochokinesis, ataxia and dysarthria and a reduction of Purkinje cells as reflected by the behavioral assays performed particularly the footprint assay and is in line with clinical observations in human LCH-ND cases.

Importantly, we found that the accumulation of circulating *BRAF*V600E^+^ cells was associated with a breakdown of the BBB with increased translocation of inflammatory cytokines and infiltration of circulating senescent *BRAF*V600E^+^ myeloid cells into the brain parenchyma. We also show that circulating *BRAF*V600E^+^ myeloid cells differentiated into senescent CD11a^+^ macrophages within the brain parenchyma. It is likely that cellular senescence of circulating *BRAF*V600E^+^ myeloid cells and brain-infiltrating *BRAF*V600E^+^ macrophages contribute to disease progression due to apoptosis resistance and production of senescent associated secretory proteins (SASP) which include inflammatory cytokines such as IL-1, IL-6, and matrix metalloproteinases (MMPs). Inflammation has long been known to disrupt the BBB and contribute to the pathogenesis of ND^24,25^. MMPs in particular have been shown to be secreted by immune cells leading to a degradation of the basement membrane and subsequent disruption of the BBB^26^. Inflammatory cytokines have also been shown to facilitate the migration of myeloid cells into the brain^27^. Importantly, pre-clinical data from LCH mouse models demonstrate that senolytic therapy (e.g., navitoclax), along with MAPK inhibition, may prevent the persistence and differentiation of senescent myeloid cells and subsequent, pathologic neuroinflammation in LCH-ND.

A recent study reported that enforced expression of *BRAF*V600E in erythromyeloid progenitors in mouse embryos leads to progressive neurodegeneration in a mouse model without systemic LCH-like disease and also identified *BRAF*V600E^+^ cells with a microglia-like phenotype in brain biopsies of patients with neurodegeneration associated with histiocytic disorders^23^. A debate has subsequently developed regarding mechanisms of LCH-ND (and ND associated with related histiocytoses such as ECD) and whether the pathogenic cells invading the brain originate from circulating *BRAF*V600E^+^ MNP that derived from *BRAF*V600E^+^ hematopoietic precursors in the bone marrow or from *BRAF*V600E^+^ microglia that seed the brain during embryonic development. Differentiating these models has significant clinical implications on surveillance and therapeutic strategies to prevent and treat LCH-ND.

A bone marrow-derived pathogenesis paradigm supports a treatment goal of molecular negativity in measurable residual disease (MRD), as is typical for malignant hematologic diseases. The ability of post-natal *BRAF*V600E^+^ myeloid precursors to recapitulate cellular and clinical features of LCH-ND – and the identification of *BRAF*V600E^+^ PBMC and the perivascular localization of *BRAF*V600E^+^ CXCR4^+^ myeloid cells in the brain of human LCH-ND – strongly support a hematopoietic origin of cellular drivers of LCH-ND. We previously reported that the extent of disease in LCH is determined by the differentiation state of the myeloid precursors in which the *BRAF*V600E mutation (or alternative activating MAPK pathway mutations) arise^10^. The patterns of disease associated with LCH-ND (e.g., disseminated LCH, lesions in skull base and facial bones, and pituitary involvement^5^) may represent *BRAF*V600E in myeloid precursors with potential to develop into both CD207^+^ DC-like cells (systemic lesions) and monocyte-derived CD11a^+^ macrophages (LCH-ND). Currently, “cure” with MRD negativity is sometimes achieved by chemotherapy but rarely achieved by MAPK inhibitor treatment alone, even in cases with positive clinical responses^22,28^. Patients with resolution of systemic lesions but persistent *BRAF*V600E^+^ peripheral blood clone could remain at risk for the development of LCH-ND. Therefore, the newly-described CD11a^+^ macrophage represents a Trojan horse, and a potential therapeutic target, responsible for trafficking LCH disease to the CNS.

The mouse models and treatment approaches developed in this study not only transform our understanding of LCH-ND pathophysiology, but also pave the way for future preventive and therapeutic treatment trajectories for LCH-ND. Beyond LCH, MAPK activation in monocyte precursors could be more broadly relevant in driving neuroinflammation in conditions such as infections or age-associated clonal hematopoiesis, where MAPK activation in monocytes from a variety of stimuli may prompt differentiation into senescent CD11a^+^ macrophages, invasion of the BBB, and neuroinflammation.

## Acknowledgements

We would like to thank Amanda Reid and Giorgio Ioannou for their excellent technical assistance. We would also like to thank the expertise and assistance of the Dean’s Flow Cytometry CORE facility, the Microscopy and Advanced Bioimaging Core facility and the Human Immune Monitoring Center (HIMC) at Mount Sinai. Further, we would like to acknowledge the Neuropathology Brain Bank & Research CoRE at Mount Sinai and especially Valeriy Borukhov for their contribution of tissue and histology services and the AP Pathology research lab at Cincinnati Children’s Hospital Medical Center for their histology assistance. This work was supported by a grant by the Swiss Cancer Research foundation (KFS-4724-02-2019 BIL) as well as by a grant from the Swiss National Science Foundation (SNSF PostDoc Mobility Fellowship P400PM_186740) to CMW. MM and CEA receive funding from the National Institute of Health (R01 CA154947), St. Baldrick’s Foundation (Consortium Grant for NACHO), the Leukemia and Lymphoma Society TRP and the HistioCure Foundation.

## Author contributions

Conceptualization, CMW, FC, OT, SAL, FG, SJR, CAE, MM

Methodology, CMW, FC, OT, PH, SB, MM

Validation, CMW, FC, OT, JLB, MDP, GRH, PH, LT, BPS, DD, AS,

Formal Analysis, CMW, FC, OT, MDP

Investigation, CMW, FC, OT, JLB, MDP, GRH, PH, LT, BPS, DD, AS, RF, MB, HL, EMT, JKK, SAL

Resources, ZY, JFC, KLM, JLP, SJR, CAE, MM

Data Curation, CMW, JLB, MDP

Writing – Original Draft, CMW, CAE, MM

Writing – Review & Editing, CMW, FC, OT, JLB, MDP, GFH, PH, BPS, MB, SB, KLM, JLP, SJR, CAE, MM

Visualization, CMW, OT, GRH, PH

Supervision, SJR, CEA, MM

Project Administration, CMW, CEA, MM Funding Acquisition, CWM, CEA, MM

## STAR Methods

### Mice

*BRAF*V600E*^HSC-Scl-CreERT^*xR26*^YFP/-^*mice (*BRAF*V600E*^Scl^* mice) and *BRAF*wt*^HSC-Scl-CreERT^*xR26*^YFP/-^*mice (*BRAF*wt*^Scl^* control mice) were generated as described previously^3^. To induce recombination in these mice, 5 doses of tamoxifen were given on 5 consecutive days (5mg, 2mg, 2mg, 1mg, 1mg) as previously described^29^. *BRAF*V600E*^Scl^*mice and *BRAF*wt*^Scl^* control mice were terminally analyzed between week 8 and 12 post tamoxifen induction. *BRAF*V600E*^Map17-CreERT^*xR26*^tdTomato/-^*mice (*BRAF*V600E*^Map17^* mice) and *BRAF*wt*^Map17-CreERT^*xR26*^tdTomato/-^*mice (*BRAF*wt*^Map17^* control mice) were generated by crossing *Map17^creER/+^R26tdTomato^+/+^* (*Pdzk1ip1-creER R26tdTomato^+/+^*)^15,20^ mice to *BRAF*V600E*^ca/WT^* mice. To induce recombination in *BRAF*V600E*^Map17^*mice and *BRAF*wt*^Map17^* control mice, 1 dose of 5mg tamoxifen was given as previously described^15,20^. *BRAF*V600E*^Map17^* mice and *BRAF*wt*^Map17^*control mice were terminally analyzed between week 12 and 16 post tamoxifen induction. BM chimera were generated by transplanting 5x10^6^ total BM cells from *BRAF*V600E*^Scl^* mice or *BRAF*wt*^Scl^*control mice into congenic CD45.1 WT host mice (*BRAF*V600E*^Scl^*chimera and *BRAF*wt*^Scl^* control chimera). Host mice were irradiated with twice 5.5Gy 6 hours apart with a lead shield protecting the head of the host mice in analogy to a previous report^16^. To induce recombination in *BRAF*V600E*^Scl^*chimera and *BRAF*wt*^Scl^* control chimera, 2 doses of tamoxifen 2mg were given 1 week apart not earlier than 4 weeks after transplantation. Chimera were terminally analyzed between week 12 and 20 after tamoxifen induction. Ethical approval for mouse experiments was obtained by the Internal Animal Care and Use Committee (IACUC) at the Mount Sinai Hospital.

### Brain macrophage isolation

Brain macrophages were isolated as described previously^14^. Mice were anesthetized with Ketamine/Xylazine and upon areflexia transcardially perfused with PBS. Brains were extracted, cut into small pieces and digested with Collagenase IV 0.2 mg/mL (Sigma, C5138-1G) and DNAse-1 0.05mg/mL (Sigma, DN25-1G) in RPMI containing 10% FCS ^30^ for 30 min at 37°C. Digested brains were passed through a 70-μm cell strainer and incubated with magnetic anti-CD45 microbeads (Miltenyi Biotec, #130-052-301) according to the manufacturer’s instructions. CD45^+^ cells were isolated using two consecutive LS columns (Miltenyi Biotec, #130-042-401) per brain and subjected to downstream analyses.

### Intravascular staining for discrimination of vascular vs parenchymal myeloid cells

Discrimination of intravascular vs extravascular myeloid cells was performed in analogy to published protocols^31,32^. Mice were anesthetized with Ketamine/Xylazine and 10μl of an anti-mouse CD45 antibody conjugated to BV510 were injected intravenously in a 100μl PBS. 10 minutes later, mice were anaesthetized and perfused with PBS. Brain macrophages were then extracted as described above and stained with respective antibodies containing an anti-mouse CD45 antibody conjugated to AF700.

### Spectral cytometry

Brain macrophage single cell suspensions were stained with a fixable blue dead cell stain kit (ThermoFisher Scientific, L-23105) in PBS for 15min on ice. After one washing cycle in cytometry buffer (PBS containing 10% BSA and 2mM EDTA), each cell sample was resuspended in 50μl of cytometry buffer containing the respective antibodies and complemented with 5μl of Super Bright Complete Staining Buffer (ThermoFisher Scientific, SB-4401-75). Samples were acquired on a Cytek® Aurora full spectrum cytometer (Configuration: 5 Laser - 16UV-16V-14B-10YG-8R). Autofluorescence detection and extraction was applied during unmixing using Cytek® SpectroFlo® v2.2.0.3 software (Cytek Biosciences, Fremont, CA, USA). Unmixed .fcs files were then analyzed with FlowJo Version 10 (Becton Dickinson, Franklin Lakes, NJ, USA). Gating strategies are displayed in **Fig. S2** and **Fig. S3**. Conditions for the respective antibodies are listed in **Table S1**.

### Cell sorting

Brain macrophage single-cell suspension and cell staining was performed as described above. Brain samples were then sorted on a Cytek® Aurora CS full spectrum cell sorter (Configuration: 5 Laser - 16UV-16V-14B-10YG-8R) with autofluorescence detection and extraction applied during unmixing (Cytek Biosciences, Fremont, CA, USA). The instrument was set up using a 100μm nozzle at 18.3psi using Single Cell sort mode. Cells were sorted into RNA lysis buffer in 1.5ml tubes. Sorted brain macrophage populations were then subjected to Ultra-low-input RNA sequencing. Conditions for the respective antibodies are listed in **Table S1**.

### Ultra-low-input RNA-sequencing

RNA extraction, library preparation, sequencing and analysis were completed at Azenta Life Science (South Plainfield, NJ) as follows:

### RNA Extraction

Total RNA was extracted using Qiagen RNeasy Plus Mini kit following manufacturer’s instructions (Qiagen, Hilden, Germany). Extracted RNA samples were quantified using Qubit 2.0 Fluorometer (Life Technologies, Carlsbad, CA, USA) and RNA integrity was checked using Agilent TapeStation 4200 (Agilent Technologies, Palo Alto, CA, USA).

### Library Preparation

Ultra-low input RNA sequencing library was prepared by using SMART-Seq HT kit for full-length cDNA synthesis and amplification (Takara, San Jose, CA, USA), and Illumina Nextera XT (Illumina, San Diego, CA, USA) library was used for sequencing library preparation. Briefly, cDNA was fragmented, and adaptor was added using Transposase, followed by limited-cycle PCR to enrich and add index to the cDNA fragments. The sequencing library was validated on the Agilent TapeStation (Agilent Technologies, Palo Alto, CA, USA), and quantified by using Qubit 2.0 Fluorometer (ThermoFisher Scientific, Waltham, MA, USA) as well as by quantitative PCR (KAPA Biosystems, Wilmington, MA, USA).

### Sequencing

The sequencing libraries were multiplexed and clustered onto a flowcell. After clustering, the flowcell was loaded onto the Illumina HiSeq instrument according to manufacturer’s instructions. The samples were sequenced using a 2x150bp Paired End (PE) configuration. Image analysis and base calling were conducted by the HiSeq Control Software (HCS). Raw sequence data (.bcl files) generated from Illumina HiSeq was converted into fastq files and de-multiplexed using Illumina bcl2fastq 2.20 software. One mis-match was allowed for index sequence identification. After investigating the quality of the raw data, sequence reads were trimmed to remove possible adapter sequences and nucleotides with poor quality using Trimmomatic v.0.36. The trimmed reads were mapped to the Mus musculus reference genome available on ENSEMBL using the STAR aligner v.2.5.2b. BAM files were generated as a result of this step. Unique gene hit counts were calculated by using feature Counts from the Subread package v.1.5.2. Only unique reads that fell within exon regions were counted. After extraction of gene hit counts, the gene hit counts table was used for downstream differential expression analysis.

### Analysis

Gene expression was computed as normalized transcript counts per million for individual samples to control for transcript sequencing variability across runs. To assess gene expression patterns across species, homologous genes between mice and humans were identified in the libraries generated for this study and for prior published work (GSE 73721 and GSE 74442)^18,33^. Respective datasets were then subsetted for these homologous genes across samples to compute log normalized gene expression as an integrated dataset. To assess transcriptomic differences driven by induction of the *BRAF*V600E mutation in hematopoietic cells, a deviance score – defined as the orthogonal distance of individual genes from the line of equality on a Cartesian plane, representative of concordance in mRNA expression – was generated to compute a quantitative description for transcriptomic remodeling.

### Immunohistochemistry and Multiplexed immunohistochemical consecutive staining on a single slide

Mice were anesthetized with Ketamine/Xylazine as previously described and upon areflexia transcardially perfused with PBS followed by 4% PFA. Brains were extracted and post-fixed in 4% PFA overnight and embedded in paraffin. Multiplexed immunohistochemical consecutive staining on a single slide (MICSSS) was performed as described previously^34^. Conditions for the respective antibodies are listed in **Table S2**.

### Luminex® multiplexed protein quantification from mouse brains

Brain tissue of perfused mice was dissociated with Pistil A and B and then lysed in ProcartaPlex™ Cell Lysis buffer (ThermoFisher Scientific, EPX-999-000) with Halt^™^ Protease Inhibitor Cocktail (ThermoFisher Scientific, #78430). Samples were spun down at 20.000xg for 10 minutes. Supernatants were frozen at -80°C until further processing. Samples were run on the FlexMap3D Luminex™ machine according to the manufacturer’s recommendations using the following Millipore panels: MKI2MAG-94K, MMMP1MAG-79K, MHSTCMAG-70KPXBK, MMMP2MAG-79K (Millipore, Burlington, MA, USA).

### Evans blue assay

*BRAF*V600E*^Scl^* mice and *BRAF*wt*^Scl^*control mice were intravenously injected with 6μl/g body weight of Evans Blue 2% in 0.9% saline. 16 hours later, animals were anesthetized with Ketamine / Xylazine and transcardially perfused with 25ml of PBS. Brains were then weighed and 400μl of dimethylformamide was added to the 1.5ml tube containing the brain and dissociated. Brains were incubated at 55°C for 48h and vortexed once daily. Samples were then spun down at 21,000g for 30min and supernatant was analyzed on a photometer with an excitation of 620nm and Emission of 680nm with an appropriate control standard curve.

### Pericyte coverage

Brain blood vessel coverage with pericytes was investigated as described previously^35^. Mice were perfused with PBS followed by 4% PFA using a gravity perfusion approach (20cm gradient and 1.2mm tube diameter). Brains were post fixed overnight in 4% PFA and then cut on a vibratome in 60μm thick sections. Free-floating tissue sections were permeabilized overnight in permeabilization buffer (1% BSA, 2% Triton-X100 in PBS) at 4°C. Primary antibodies directed against Collagen IV (Bio-Rad, #2150-1470, 1:300) and CD13 (R&D Systems, AF2335, 1:100) were incubated for 48 hours followed by secondary antibodies (Jackson Immuno Research Cy5™-conjugated AffiniPure Donkey Anti-Goat IgG #705-175-147 and DyLight™488 AffiniPure Donkey Anti-Rabbit IgG #711-485-152, dilution 1:300) for 24 hours. Tissue sections were mounted using Molecular Probes ProLong Gold Antifade Mountant (ThermoFisher Scientific, #P36930). Conditions for the respective antibodies are also listed in **Table S3** and **4**.

### Quantification of vessel pericyte coverage

Images of immune-fluorescently labeled sections were acquired by Zeiss LSM780 confocal laser scanning microscope with a 20X objective (Zeiss, Oberkochen, Germany). Pericyte coverage was calculated using the area measurement tool in Fiji. The area of CD13 and Collagen IV signal was measured on binary images in 6 ROIs, 100 x 100 µm each, in sagittal brain section. Coverage was calculated as the percentage of CD13 positive area over the Collagen IV positive area. The Collagen IV area was taken arbitrarily as 100% and the CD13 positive area was expressed as a percentage normalized to the Collagen IV area.

### Biotinylation

Biotinylation of recombinant IL-1β (R&D Systems, 201-LB-025/CF) was performed using the EZ-Link™ Sulfo-NHS-Biotin kit according to the manufacturer’s instructions (ThermoFisher Scientific, #A39256). Biotinylated IL-1β was separated from unbound biotin using Pierce™ C18 Spin Columns, 7K MWCO, (ThermoFisher Scientific, #89870), which recovers proteins and macromolecules larger than 7kDa. 100μl of biotinylated IL-1β was injected retro-orbitally into anaesthetized mice at a concentration of 250ng/ml. After 2 hours of circulation, mice were euthanized and perfused with ice cold PBS followed by 4% PFA. Brain tissue processing and imaging was performed as described in the Pericyte coverage section. Biotin was visualized using the Oregon Green® 488 conjugate of NeutrAvidin® biotin-binding protein (ThermoFisher Scientific, #A6374). Counterstaining was performed using rabbit anti-Collagen IV (1:300, Bio-Rad, #2150-1470). Conditions for the respective antibodies are also listed in **Table S3** and **4**.

### Open Field Assay

*BRAF*V600E*^Scl^* chimera and *BRAF*wt*^Scl^* chimera were studied for abnormal behavior in an Open Field assay. Therefore, movement of mice was tracked in an open field chamber for 60 minutes during the night cycle after acclimatization to red light for one hour. Data were analyzed using the proprietary software of the Open Field chambers (Omnitech Electronics, Fusion 5.6).

### Grip strength analysis

*BRAF*V600E*^Scl^* chimera and *BRAF*wt*^Scl^*control chimera were tested for grip strength using a grip strength meter (Biosep, Bio-GS3) as suggested by the manufacturer. Mice were trained on the setup before grip strength was assess in 3 consecutive rounds including appropriate breaks between sessions. Mean values of the peak grip force of each mouse were used for statistical analysis.

### Rotarod Assay

*BRAF*V600E*^Scl^* chimera and *BRAF*wt*^Scl^*chimera were evaluated for ataxia using an accelerating rotating rod (Rotarod) setup^36^. The animals were placed on the rotating rod an acceleration from 4 to 40rpm was initiated. Latency to fall from the rod was recorded after training runs. In total 3 recorded sessions were analyzed and mean latency to fall plotted.

### Footprint analysis

*BRAF*V600E*^Scl^* chimera and *BRAF*wt*^Scl^*control chimera were investigated for cerebellar ataxia using a footprint assay as previously described^37^.

### *In vivo* drug treatment

*BRAF*V600E*^Scl^*mice and *BRAF*wt*^Scl^*control mice as well as *BRAF*V600E*^Scl^* chimera and *BRAF*wt*^Scl^*control chimera were treated with Navitoclax at a dose of 50mg/kg daily by oral gavage (ApexBio, #A3007)^3^ in combination with Trametinib 1mg/kg daily intraperitoneally (i.p.) (Selleck Chemicals, #S2673)^38^ for 8 weeks before terminal analysis.

### Data availability

Datasets supporting the findings presented in this study are available from the corresponding author upon reasonable request. Any data that can be shared will be released via a material transfer agreement. Bulk RNA-seq data obtained from mouse brain macrophages will be made available upon request.

### Statistical analysis

Statistical significance between groups was determined by unpaired Student’s t-test or one-sided ANOVA; data are displayed as means ± s.e.m. Statistical analysis was done with GraphPad Prism v.9.3.

---

**Figure S1:**
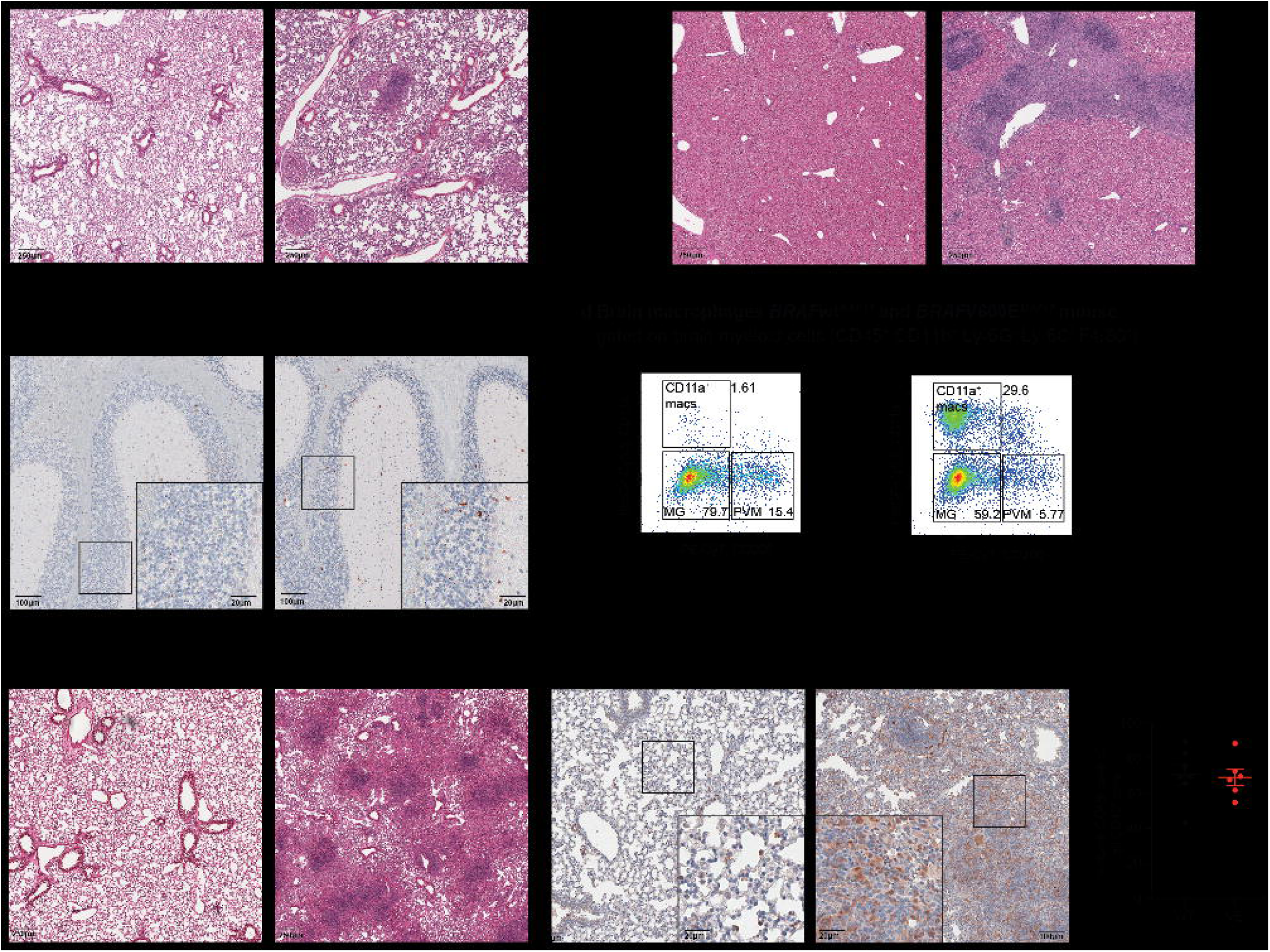
LCH phenotype of *BRAF*V600E*^Map17^*mice and *BRAF*V600E*^Scl^* chimera in comparison to respective control mice. **a,** representative H&E images of lung tissue from a *BRAF*V600E*^Map17^* (right) mouse and a *BRAF*wt*^Map17^*control mouse (left) showing typical granulomatous lesions in this LCH mouse model 12-16 weeks post Tamoxifen injection **b**, representative H&E images of liver tissue of these mice showing immune cell infiltrates in *BRAF*V600E*^Map17^* (right) mice but not in *BRAF*wt*^Map17^*control mice (left). **c**, representative IHC staining of the *BRAF*V600E reporter tdTomato in the brain tissue of *BRAF*V600E*^Map17^* revealing a diffuse infiltration of the brains of *BRAF*V600E*^Map17^* mice (right panel) but not *BRAF*wt*^Map17^* control mice 12-16 weeks post Tamoxifen injection (left panel). **d**, representative spectral cytometry pseudocolor plots of brain microglia (MG), perivascular macrophages (PVM) and CD11a^+^ macrophages of *BRAF*V600E*^Map17^* mice and *BRAF*wt*^Map17^*control mice confirming the abundance of the pathognomonic CD11a^+^ macrophage population in the brains of *BRAF*V600E*^Map17^* mice. **e**, representative H&E stain of lungs from *BRAF*V600E*^Scl^*chimera (right) and *BRAF*wt*^Scl^* control chimera (left) showing typical granulomatous LCH-like lesions in *BRAF*V600E*^Scl^*chimera. **f**, representative IHC staining of the reporter protein YFP in the lungs of *BRAF*V600E*^Scl^* chimera demonstrating that *BRAF*V600E-mutated cells accumulate in granulomatous lesions in the lungs of *BRAF*V600E*^Scl^* chimera (right) while scattered YFP-tagged cells can be detected in the lungs of *BRAF*wt*^Scl^*control chimera (left). **g**, donor chimerism in *BRAF*V600E*^Scl^*chimera and *BRAF*wt*^Scl^* control chimera at terminal analysis (CD45.2 as percent of total CD45) showing comparable chimerism between *BRAF*V600E*^Scl^* chimera and *BRAF*wt*^Scl^*control chimera.

**Figure S2:**
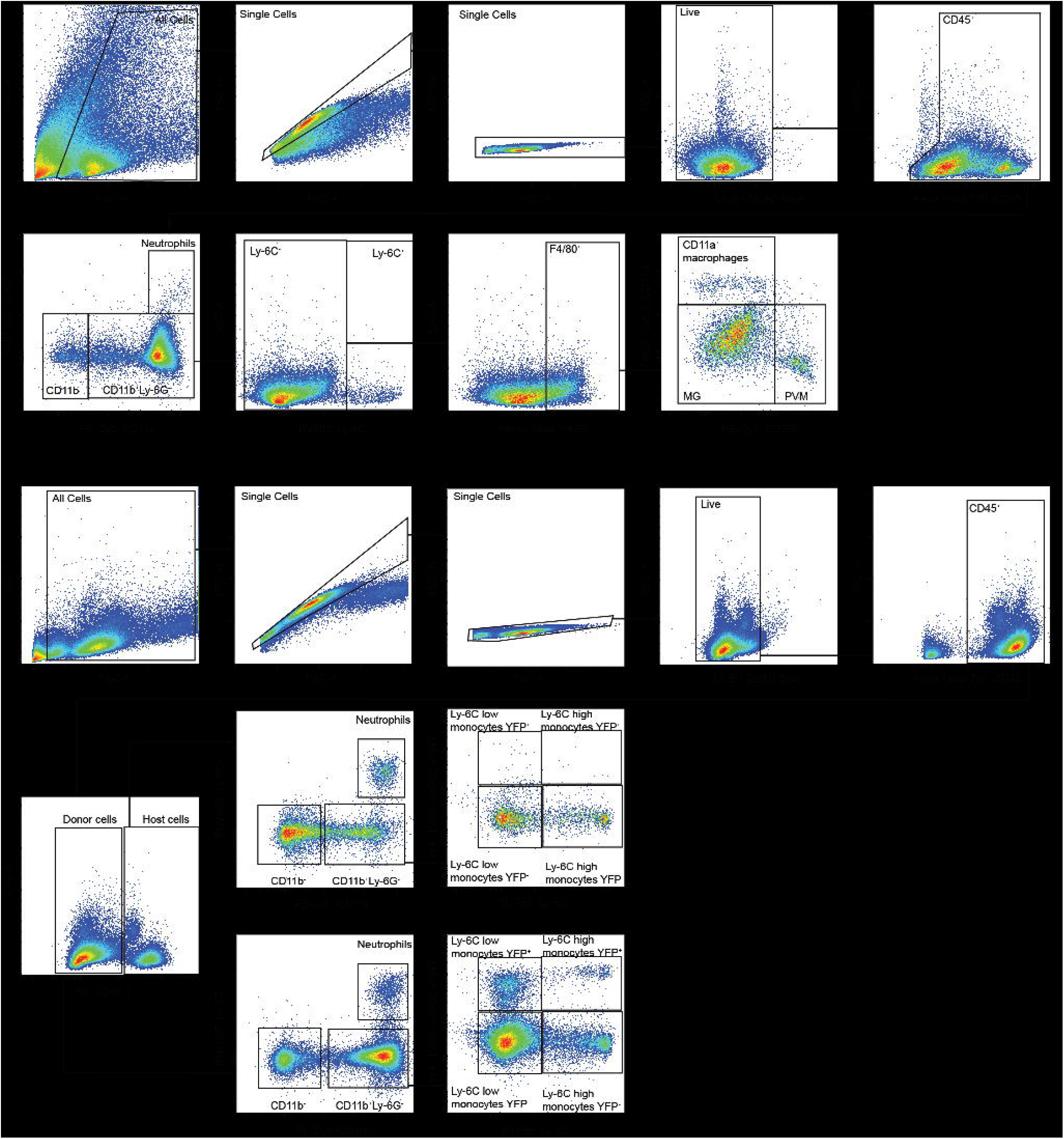
Gating Strategies. **a,** gating strategy for brain myeloid cells from *BRAF*V600E*^Scl^* mice and *BRAF*wt*^Scl^*control mice and *BRAF*V600E*^Scl^* chimera and *BRAF*wt*^Scl^*control chimera. **b,** gating strategy for peripheral blood cells in *BRAF*V600E*^Scl^* chimera and *BRAF*wt*^Scl^* control chimera.

**Figure S3:**
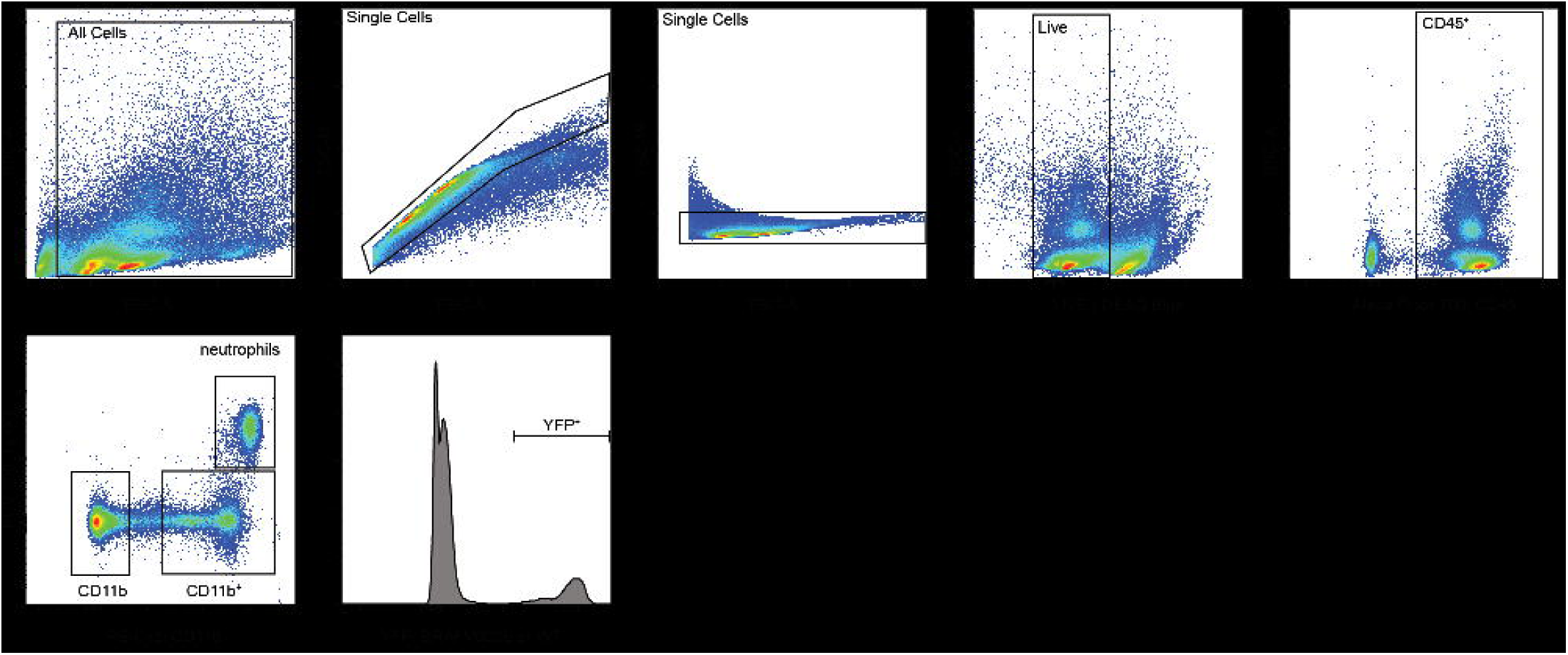
Gating Strategies. **a**, gating strategy for lung immune cells from *BRAF*V600E*^Scl^* mice and *BRAF*wt*^Scl^* control mice.

**Table S1.** Antibodies used for Spectral Cytometry.

| Antigen | Reactivity | Clone | Host | Conjugate | Company | Order number |
| --- | --- | --- | --- | --- | --- | --- |
| CD4 | mouse | rat | GK1.5 | BV750 | biolegend | 100467 |
| CD11b | mouse/human | rat | M1/70 | CD11b | biolegend | 101209 |
| CD45 | mouse | rat | 30-F11 | AF700 | biolegend | 103127 |
| CD45 | mouse | rat | 30-F11 | BV510 | biolegend | 103137 |
| CD45.1 | mouse | mouse | A20 | PE | biolegend | 110707 |
| MHCII | mouse | rat | M5/114.15.2 | BV711 | biolegend | 107643 |
| Ly-6C | mouse | rat | HK1.4 | BV785 | biolegend | 128041 |
| CD8a | mouse | mouse | QA17A07 | BV421 | biolegend | 155010 |
| Ly-6G | mouse | rat | 1A8 | BUB805 | BD Biosciences | 741994 |
| B220 | mouse | rat | RA3-6B2 | BV570 | biolegend | 103237 |
| CD11a | mouse | rat | M17/4 | PerCP/Cy5.5 | biolegend | 101123 |
| CD206 | mouse | rat | C068C2 | PE/Cy7 | biolegend | 141720 |
| F4/80 | mouse | rat | BM8 | efluor450 | ThermoFisher | 48-4801-82 |
| F4/80 | mouse | rat | BM8 | APC-Fire750 | biolegend | 123151 |
| F4/80 | mouse | rat | BM8 | BV605 | biolegend | 123133 |

**Table S2.** Antibodies used for Immunohistochemistry.

| Antigen | Reac-tivity | Clone | Host | AG retr | Dilution | Incu-bation | Company | Order number |
| --- | --- | --- | --- | --- | --- | --- | --- | --- |
| Cal-bindin-1 | mouse | Poly-clonal | Mouse | pH 9 | 1500 | 1h RT | Proteintech | 14479-1 |
| YFP GFP | n/a | JL8 | Mouse | pH 9 | 200 | 4°C o/n | Takara Bio | 632381 |
| tdTomato RFP | n/a | Poly-clonal | Mouse | pH 9 | 200 | 1h RT | Takara Bio | 632496 |
| GFAP | mouse/hu | Poly-clonal | Rabbit | pH 9 | 300 | 1h RT | Agilent | Z033429-2 |

**Table S3.** Antibodies used for Immunoflourescence.

| Antigen | Reactivity | Clone | Host | Dilution | Incu-bation | Company | Order number |
| --- | --- | --- | --- | --- | --- | --- | --- |
| Aminopeptidase N / CD13 | Mouse | Poly-clonal | goat | 100 | 48h | R&D Systems | AF2335 |
| Collagen IV | Mouse | Poly-clonal | rabbit | 300 | 48h | Bio-Rad | 2150-1470 |

**Table S4.** secondary antibodies / dyes Immunofluorescence.

| reactivity | clone | host | Con-<br>jugate | Dilution | Incu-bation | company | order no |
| --- | --- | --- | --- | --- | --- | --- | --- |
| goat | Poly-clonal | Donkey | Cy5 | 400 | 24h | Jackson<br>Immuno<br>Research | #705-175-147 |
| rabbit | Poly-clonal | Donkey | AF488 | 400 | 24h | Jackson<br>Immuno<br>Research | #711-485-152 |
| Biotin | n/a | n/a | AF488 | 300 | 24h | Thermofisher | #A6374 |

## References

1. Allen, C.E., Flores, R., Rauch, R., Dauser, R., Murray, J.C., Puccetti, D., Hsu, D.A., Sondel, P., Hetherington, M., Goldman, S., et al. (2010). Neurodegenerative central nervous system Langerhans cell histiocytosis and coincident hydrocephalus treated with vincristine/cytosine arabinoside. Pediatr Blood Cancer 54, 416–423. 10.1002/pbc.22326.

2. Héritier, S., Barkaoui, M., Miron, J., Thomas, C., Moshous, D., Lambilliotte, A., Mazingue, F., Kebaili, K., Jeziorski, E., Plat, G., et al. (2018). Incidence and risk factors for clinical neurodegenerative Langerhans cell histiocytosis: a longitudinal cohort study. Brit J Haematol 183, 608–617. 10.1111/bjh.15577.

3. Bigenwald, C., Berichel, J.L., Wilk, C.M., Chakraborty, R., Chen, S.T., Tabachnikova, A., Mancusi, R., Abhyankar, H., Casanova-Acebes, M., Laface, I., et al. (2021). BRAFV600E-induced senescence drives Langerhans cell histiocytosis pathophysiology. Nat Med, 1–11. 10.1038/s41591-021-01304-x.

4. Allen, C.E., Merad, M., and McClain, K.L. (2018). Langerhans-Cell Histiocytosis. New Engl J Medicine 379, 856–868. 10.1056/nejmra1607548.

5. Yeh, E.A., Greenberg, J., Abla, O., Longoni, G., Diamond, E., Hermiston, M., Tran, B., Rodriguez-Galindo, C., Allen, C.E., McClain, K.L., et al. (2018). Evaluation and treatment of Langerhans cell histiocytosis patients with central nervous system abnormalities: Current views and new vistas. Pediatr Blood Cancer 65, e26784. 10.1002/pbc.26784.

6. Grois, N., Prayer, D., Prosch, H., and Lassmann, H. (2005). Neuropathology of CNS disease in Langerhans cell histiocytosis. Brain 128, 829–838. 10.1093/brain/awh403.

7. Grois, N., Tsunematsu, Y., Barkovich, A.J., and Favara, B.E. (1994). Central nervous system disease in Langerhans cell histiocytosis. Br J Cancer Suppl 23, S24–8.

8. Idbaih, A., Donadieu, J., Barthez, M.A., Geissmann, F., Bertrand, Y., Hermine, O., Brugières, L., Genereau, T., Thomas, C., and HoanglJXuan, K. (2004). Retinoic acid therapy in “degenerativelJlike” neurolJlangerhans cell histiocytosis: A prospective pilot study. Pediatr Blood Cancer 43, 55–58. 10.1002/pbc.20040.

9. (JLSG), J.L.S.G., Imashuku, S., Fujita, N., Shioda, Y., Noma, H., Seto, S., Minato, T., Sakashita, K., Ito, N., Kobayashi, R., et al. (2015). Follow-up of pediatric patients treated by IVIG for Langerhans cell histiocytosis (LCH)-related neurodegenerative CNS disease. Int J Hematol 101, 191–197. 10.1007/s12185-014-1717-5.

10. Berres, M.-L., Lim, K.P.H., Peters, T., Price, J., Takizawa, H., Salmon, H., Idoyaga, J., Ruzo, A., Lupo, P.J., Hicks, M.J., et al. (2014). BRAF-V600E expression in precursor versus differentiated dendritic cells defines clinically distinct LCH risk groupsLCH: neoplasia arising from myeloid precursors. J Exp Medicine 211, 669–683. 10.1084/jem.20130977.

11. Hogstad, B., Berres, M.-L., Chakraborty, R., Tang, J., Bigenwald, C., Serasinghe, M., Lim, K.P.H., Lin, H., Man, T.-K., Remark, R., et al. (2018). RAF/MEK/extracellular signal–related kinase pathway suppresses dendritic cell migration and traps dendritic cells in Langerhans cell histiocytosis lesions. J Exp Med 215, 319–336. 10.1084/jem.20161881.

12. McClain, K.L., Bigenwald, C., Collin, M., Haroche, J., Marsh, R.A., Merad, M., Picarsic, J., Ribeiro, K.B., and Allen, C.E. (2021). Histiocytic disorders. Nat Rev Dis Primers 7, 73. 10.1038/s41572-021-00307-9.

13. Paolicelli, R.C., Sierra, A., Stevens, B., Tremblay, M.-E., Aguzzi, A., Ajami, B., Amit, I., Audinat, E., Bechmann, I., Bennett, M., et al. (2022). Microglia states and nomenclature: A field at its crossroads. Neuron 110, 3458–3483. 10.1016/j.neuron.2022.10.020.

14. Silvin, A., Uderhardt, S., Piot, C., Mesquita, S.D., Yang, K., Geirsdottir, L., Mulder, K., Eyal, D., Liu, Z., Bridlance, C., et al. (2022). Dual ontogeny of disease-associated microglia and disease inflammatory macrophages in aging and neurodegeneration. Immunity. 10.1016/j.immuni.2022.07.004.

15. Sawai, C.M., Babovic, S., Upadhaya, S., Knapp, D.J.H.F., Lavin, Y., Lau, C.M., Goloborodko, A., Feng, J., Fujisaki, J., Ding, L., et al. (2016). Hematopoietic Stem Cells Are the Major Source of Multilineage Hematopoiesis in Adult Animals. Immunity 45, 597–609. 10.1016/j.immuni.2016.08.007.

16. Mildner, A., Schmidt, H., Nitsche, M., Merkler, D., Hanisch, U.-K., Mack, M., Heikenwalder, M., Brück, W., Priller, J., and Prinz, M. (2007). Microglia in the adult brain arise from Ly-6ChiCCR2+ monocytes only under defined host conditions. Nat Neurosci 10, 1544–1553. 10.1038/nn2015.

17. McClain, K.L., Picarsic, J., Chakraborty, R., Zinn, D., Lin, H., Abhyankar, H., Scull, B., Shih, A., Lim, K.P.H., Eckstein, O., et al. (2018). CNS Langerhans cell histiocytosis: Common hematopoietic origin for LCH-associated neurodegeneration and mass lesions: Hematopoietic Origin of LCH-ND. Cancer 124, 2607– 2620. 10.1002/cncr.31348.

18. Diamond, E.L., Durham, B.H., Haroche, J., Yao, Z., Ma, J., Parikh, S.A., Wang, Z., Choi, J., Kim, E., Cohen-Aubart, F., et al. (2016). Diverse and Targetable Kinase Alterations Drive Histiocytic Neoplasms. Cancer Discov 6, 154–165. 10.1158/2159-8290.cd-15-0913.

19. Maier, B., Leader, A.M., Chen, S.T., Tung, N., Chang, C., LeBerichel, J., Chudnovskiy, A., Maskey, S., Walker, L., Finnigan, J.P., et al. (2020). A conserved dendritic-cell regulatory program limits antitumour immunity. Nature 580, 257–262. 10.1038/s41586-020-2134-y.

20. Casanova-Acebes, M., Dalla, E., Leader, A.M., LeBerichel, J., Nikolic, J., Morales, B.M., Brown, M., Chang, C., Troncoso, L., Chen, S.T., et al. (2021). Tissue-resident macrophages provide a pro-tumorigenic niche to early NSCLC cells. Nature, 1–7. 10.1038/s41586-021-03651-8.

21. Aubart, F.C., Emile, J.-F., Carrat, F., Charlotte, F., Benameur, N., Donadieu, J., Maksud, P., Idbaih, A., Barete, S., Hoang-Xuan, K., et al. (2017). Targeted therapies in 54 patients with Erdheim-Chester disease, including follow-up after interruption (the LOVE study). Blood 130, 1377–1380. 10.1182/blood-2017-03-771873.

22. Eckstein, O.S., Visser, J., Rodriguez-Galindo, C., Allen, C.E., and Group, N.-L.S. (2019). Clinical responses and persistent BRAF V600E+ blood cells in children with LCH treated with MAPK pathway inhibition. Blood 133, 1691–1694. 10.1182/blood-2018-10-878363.

23. Mass, E., Jacome-Galarza, C.E., Blank, T., Lazarov, T., Durham, B.H., Ozkaya, N., Pastore, A., Schwabenland, M., Chung, Y.R., Rosenblum, M.K., et al. (2017). A somatic mutation in erythro-myeloid progenitors causes neurodegenerative disease. Nature 549, 389–393. 10.1038/nature23672.

24. Varatharaj, A., and Galea, I. (2017). The blood-brain barrier in systemic inflammation. Brain Behav Immun 60, 1–12. 10.1016/j.bbi.2016.03.010.

25. Takata, F., Nakagawa, S., Matsumoto, J., and Dohgu, S. (2021). Blood-Brain Barrier Dysfunction Amplifies the Development of Neuroinflammation: Understanding of Cellular Events in Brain Microvascular Endothelial Cells for Prevention and Treatment of BBB Dysfunction. Front Cell Neurosci 15, 661838. 10.3389/fncel.2021.661838.

26. Lakhan, S.E., Kirchgessner, A., Tepper, D., and Leonard, A. (2013). Matrix Metalloproteinases and Blood-Brain Barrier Disruption in Acute Ischemic Stroke. Front Neurol 4, 32. 10.3389/fneur.2013.00032.

27. Spath, S., Komuczki, J., Hermann, M., Pelczar, P., Mair, F., Schreiner, B., and Becher, B. (2017). Dysregulation of the Cytokine GM-CSF Induces Spontaneous Phagocyte Invasion and Immunopathology in the Central Nervous System. Immunity 46, 245–260. 10.1016/j.immuni.2017.01.007.

28. Donadieu, J., Larabi, I.A., Tardieu, M., Visser, J., Hutter, C., Sieni, E., Kabbara, N., Barkaoui, M., Miron, J., Chalard, F., et al. (2019). Vemurafenib for Refractory Multisystem Langerhans Cell Histiocytosis in Children: An International Observational Study. J Clin Oncol 37, 2857–2865. 10.1200/jco.19.00456.

29. Göthert, J.R., Gustin, S.E., Hall, M.A., Green, A.R., Göttgens, B., Izon, D.J., and Begley, C.G. (2005). In vivo fate-tracing studies using the Scl stem cell enhancer: embryonic hematopoietic stem cells significantly contribute to adult hematopoiesis. Blood 105, 2724–2732. 10.1182/blood-2004-08-3037.

30. Liu, Z., Gu, Y., Shin, A., Zhang, S., and Ginhoux, F. (2020). Analysis of Myeloid Cells in Mouse Tissues with Flow Cytometry. Star Protoc 1, 100029. 10.1016/j.xpro.2020.100029.

31. Anderson, K.G., Mayer-Barber, K., Sung, H., Beura, L., James, B.R., Taylor, J.J., Qunaj, L., Griffith, T.S., Vezys, V., Barber, D.L., et al. (2014). Intravascular staining for discrimination of vascular and tissue leukocytes. Nat Protoc 9, 209–222. 10.1038/nprot.2014.005.

32. Hamon, P., Loyher, P.-L., Chanville, C.B. de, Licata, F., Combadière, C., and Boissonnas, A. (2017). CX3CR1-dependent endothelial margination modulates Ly6Chigh monocyte systemic deployment upon inflammation in mice. Blood 129, 1296–1307. 10.1182/blood-2016-08-732164.

33. Zhang, Y., Sloan, S.A., Clarke, L.E., Caneda, C., Plaza, C.A., Blumenthal, P.D., Vogel, H., Steinberg, G.K., Edwards, M.S.B., Li, G., et al. (2016). Purification and Characterization of Progenitor and Mature Human Astrocytes Reveals Transcriptional and Functional Differences with Mouse. Neuron 89, 37–53. 10.1016/j.neuron.2015.11.013.

34. Remark, R., Merghoub, T., Grabe, N., Litjens, G., Damotte, D., Wolchok, J.D., Merad, M., and Gnjatic, S. (2016). In-depth tissue profiling using multiplexed immunohistochemical consecutive staining on single slide. Sci Immunol 1, aaf6925. 10.1126/sciimmunol.aaf6925.

35. Török, O., Schreiner, B., Schaffenrath, J., Tsai, H.-C., Maheshwari, U., Stifter, S.A., Welsh, C., Amorim, A., Sridhar, S., Utz, S.G., et al. (2021). Pericytes regulate vascular immune homeostasis in the CNS. Proc National Acad Sci 118. 10.1073/pnas.2016587118.

36. Brooks, S.P., and Dunnett, S.B. (2009). Tests to assess motor phenotype in mice: a user’s guide. Nat Rev Neurosci 10, 519–529. 10.1038/nrn2652.

37. Sugimoto, H., and Kawakami, K. (2019). Low-cost Protocol of Footprint Analysis and Hanging Box Test for Mice Applied the Chronic Restraint Stress. J Vis Exp. 10.3791/59027.

38. Sengal, A., Velazquez, J., Hahne, M., Burke, T.M., Abhyankar, H., Reyes, R., Olea, W., Scull, B., Eckstein, O.S., Bigenwald, C., et al. (2021). Overcoming T-cell exhaustion in LCH: PD-1 blockade and targeted MAPK inhibition are synergistic in a mouse model of LCH. Blood 137, 1777–1791. 10.1182/blood.2020005867.

